# PCR Bias Impacts Microbiome Ecological Analyses

**DOI:** 10.1101/2025.07.31.667904

**Authors:** Dharmik Rathod, Justin D. Silverman

**Affiliations:** College of Information Sciences and Technology, The Pennsylvania State University; Department of Statistics, The Pennsylvania State University; Department of Medicine, The Pennsylvania State University; Institute for Computational and Data Science, The Pennsylvania State University

## Abstract

Polymerase Chain Reaction (PCR) is a critical step in amplicon-based microbial community profiling, allowing the selective amplification of marker genes such as 16S rRNA from environmental or host-associated samples. Despite its widespread use, PCR is known to introduce amplification bias, where some DNA sequences are preferentially amplified over others due to factors such as primer-template mismatches, sequence GC content, and secondary structures. Although these biases are known to affect transcript abundance, their implications for ecological metrics remain poorly understood. In this study, we conduct a comprehensive evaluation of how PCR-bias influences both within-samples (*α*-diversity) and between-sample (*β*-diversity) analyses. We show that perturbation-invariant diversity measures remain unaffected by PCR bias, but widely used metrics such as Shannon diversity and Weighted-Unifrac are sensitive, with their values varying according to the true community composition. To address this, we provide theoretical and empirical insight into how PCR-induced bias varies across ecological analyses and community structures, and we offer practical guidance on when bias-correction methods should be applied. Our findings highlight the importance of selecting appropriate diversity metrics for PCR-based microbial ecology workflows and offer guidance for improving the reliability of diversity analyses.

## 1 Introduction

The emergence of next-generation sequencing technologies, along with advances in computational tools, has substantially expanded microbiome research across biomedicine [Tringe and Hugenholtz, 2008]. Amplicon-based microbiome studies typically target hypervariable regions of the 16S rRNA gene, using them as barcodes to quantify the relative abundance of bacterial taxa in mixed communities [Youssef et al., 2009, Caporaso et al., 2011]. To isolate the 16S rRNA gene and generate sufficient material for sequencing, multiple cycles of polymerase chain reaction (PCR) amplification are usually required [Pinto and Bhatt, 2024]. However, amplification efficiency can vary across 16S rRNA templates, leading to systematic over- or under-representation of certain taxa in sequencing libraries [Acinas et al., 2005, Suzuki and Giovannoni, 1996, Polz and Cavanaugh, 1998]. These biases can lead to over 30-fold errors in observed relative abundances, severely distorting estimates of microbial composition [Silverman et al., 2021]. Overall, PCR bias can be a substantial source of error in 16S rRNA studies [Acinas et al., 2005, Eisenstein, 2018, Polz and Cavanaugh, 1998, Suzuki and Giovannoni, 1996, McLaren et al., 2019, Silverman et al., 2021, Korvigo et al., 2022]. Despite efforts to optimize experimental protocols (e.g., Krehenwinkel et al. [2017]), PCR bias remains an outstanding problem.

PCR bias likely originates from multiple distinct processes. Mismatches between primer and template sequences are one source of bias [Parada et al., 2016]. Yet, this primer-mismatch bias is unlikely to persist past the initial cycles of PCR, after which point the template sequence is replaced by a sequence complementary to the primers themselves [Wu et al., 2009]. Still other transcript level factors can lead to Non-Primer-Mismatch (NPM) bias which persist through later cycles [Silverman et al., 2021]. For example, GC-rich templates are more stable and can require higher melting temperatures or can form hairpin structures that hamper amplification [Frey et al., 2008]. Mock community studies suggest that sources of NPM bias dominates primer-mismatch biases [Silverman et al., 2021, Korvigo et al., 2022, Gimpel et al., 2024]. Because NPM bias is the dominant and persistent source of PCR distortion, we use the terms “PCR bias” and “NPM bias” interchangeably throughout this work.

Recent studies have shown that PCR bias is highly consistent across PCR cycles and can be accurately modeled by a simple exponential bias model. Let *a*_1_*/a*_2_ represent the true ratio of two templates before amplification (i.e., at cycle 0), and let *b*_1_*/b*_2_ denote the ratio of their per-cycle amplification efficiencies, where each *b*_*j*_ ∈ [1, 2], with 1 indicating no amplification and 2 indicating perfect doubling. After *x*_*n*_ cycles of PCR, the observed ratio of the two templates becomes *w*_1*n*_*/w*_1*n*_. These quantities are related through the following model:

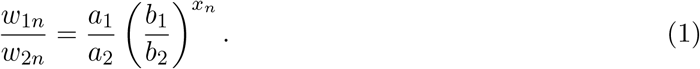

This model was originally proposed by Suzuki and Giovannoni [1996] who used it to study PCR reactions with two templates. This model was recently generalized by Silverman et al. [2021] and McLaren et al. [2019] who extended it to the case where there are more than two templates. Silverman et al. [2021] further generalized this model to situations where the ratios (e.g.,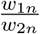) are not measured directly but observed only through noisy sequencing data. By sequencing 16S rRNA libraries with different numbers of PCR cycles, those authors could directly infer relative PCR efficiencies (e.g., 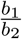) and the unbiased ratio abundances (e.g., 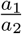). They found that this simple model explained over 95% of variation due to PCR bias in both mock and real 16S rRNA libraries. Those findings have subsequently been validated by multiple groups [Gimpel et al., 2024, Korvigo et al., 2022].

Despite advances in modeling PCR bias, relatively little research has investigated which types of microbiome analyses are most affected. For instance, it remains unclear when researchers should perform calibration experiments, such as sequencing replicates with varying numbers of PCR cycles, to enable application of models like Equation (1). Most existing studies have focused on how PCR bias distorts estimates of relative abundance, leading to systematic over- or under-representation of certain taxa. However, such distortions may not affect all downstream analyses equally. For example, both Silverman et al. [2021] and McLaren et al. [2019] have shown that differential abundance analyses, which aim to estimate changes in relative abundance across conditions (e.g., health vs. disease), can be invariant to PCR bias. In contrast, other commonly used approaches remain under-explored. In particular, ecological diversity measures such as *α*-diversity (e.g., Shannon index) and *β*-diversity (e.g., Bray–Curtis dissimilarity) form the basis of many microbiome analyses, yet the extent to which PCR bias affects these metrics has not been systematically studied.

This article presents the first systematic evaluation of how PCR bias affects ecological analyses based on *α*- and *β*-diversity metrics. First, we prove that there exists a class of perturbation-invariant diversity measures that are unaffected by PCR bias. Second, we demonstrate that a broader class of perturbation-sensitive metrics—including Shannon diversity and Bray–Curtis dissimilarity—are impacted by PCR bias. Importantly, this bias is not consistent: it depends on the true underlying composition (e.g., 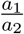), implying that PCR bias can also distort differential diversity analyses aimed at identifying changes in diversity across groups. To guide future research, we provide intuitive explanations of how bias varies with community composition and which types of microbial communities are most susceptible. Based on our findings, we offer a simple recommendation: researchers who adopt perturbation-invariant metrics need not perform calibration experiments to mitigate PCR bias, whereas those using perturbation-sensitive metrics should strongly consider doing so.

## 2. Results

### 2.1 A Statistical Model for PCR Bias

This article uses the Silverman et al. [2021] model which extends Equation (1): accounting for more than two taxa and measurement error. Note that Equation (1) can be written as a linear log-contrast model:

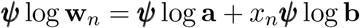

***ψ*** = (1, −1), **w**_*n*_ = (*w*_1*n*_, *w*_2*n*_)^*T*^, **a** = (*a*_1_, *a*_2_)^*T*^ and **b** = (*b*_1_, *b*_2_)^*T*^ are 2-vectors. The vector ***ψ*** is an example of a contrast, a vector with elements −1 ≤ *ψ*_*d*_ ≤ 1 that sums to 0. To extend this model beyond *D* = 2 taxa we use this linear log-contrast form.

The vector ***ψ*** can be thought of as a 1 × 2-matrix. In moving to *D >* 2 we now consider a (*D* − 1) × *D* contrast matrix **Ψ** with columns that sum to zero: 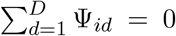 for all *i* ∈ {1, …, *D* − 1}. We now have a multivariate linear log-contrast model:

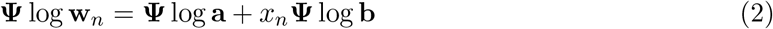

where **w, a**, and **b** are all *D*-vectors. For our purposes, we can choose any contrast matrix **Ψ** with rank *D* − 1, e.g., a matrix **Ψ** = (**I**_*D−*1_; −**1**_*D−*1_) which corresponds to the contrast matrix of the Additive Log-Ratio Transform [Pawlowsky-Glahn et al., 2015]. We use the following notation as a shorthand: ***η***_*i*_ = **Ψ** log **w**_*i*_, ***α*** = **Ψ** log **a**, and ***β*** = **Ψ** log **b** resulting in a linear model: ***η***_*n*_ = ***α*** + ***β****x*_*n*_.

Sequencing itself is a noisy measurement process that produces a zero-ladden *D*×*N* -dimensional count matrix ***Y*** with element ***Y***_*dn*_ representing the number of sequencing reads mapping to taxon *d* in sample *n*. As a result, we cannot measure ***η*** directly, or calculate it directly from ***Y***, instead, Silverman et al. [2021] proposed the following Bayesian hierarchical model which treats ***η*** as nuissance parameters to be estimated along with the parameters of interest ***α*** and ***β***:

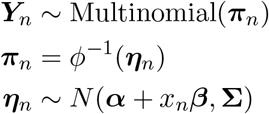

where ***π***_*n*_ is a *D*-dimensional vector of relative abundances, *ϕ*(***π***_*n*_) = **Ψ** log ***π***_*n*_ is a (*D* − 1) dimensional vector of log-ratios and **Σ** is a (*D* − 1)-dimensional covariance matrix that is also estimated from the observed data. As in Silverman et al. [2021], we estimate the parameters in this model (***α*** and ***β***) and the nuisance parameters (***η*** and **Σ**) using the *fido* R library Silverman et al. [2022]. This model has been experimentally validated by multiple groups [Korvigo et al., 2022, Gimpel et al., 2024, Silverman et al., 2021].

When applied to datasets that include calibration samples—i.e., samples sequenced after undergoing different numbers of PCR cycles—this model enables inference of the taxon-specific relative amplification efficiencies ***β*** and the underlying unbiased relative abundances ***α***, with quantified uncertainty. In practice, we often include additional covariates (beyond cycle number, *x*_*n*_) to account for batch effects. By including additional covariates, this model can also be applied to samples from distinct microbial communities, each with their own unbiased relative abundance vector *α* (See Methods 4.1).

To illustrate the model, we reproduce the analysis of Silverman et al. [2021] to estimate how PCR bias alters relative abundance estimates in both an *in vitro* and an *ex vivo* human gut microbiome study. Supplemental Figure S1, which parallels figures from the original study, visually demonstrates the extent of this distortion. Briefly, some taxa, such as Holdemania, are consistently underrepresented by a factor of approximately 32, whereas others, such as Bacteroides, are over-represented by a factor of about 4. Overall, many taxa show clear evidence of PCR bias, with 95% credible intervals for their amplification efficiencies excluding the null (no bias).

### 2.2 Sensitivity and Invariance of Estimands to PCR Bias

Most prior work on PCR bias has focused on its effect on relative abundance estimates (e.g., Silverman et al. [2021]). However, relative abundances are often not the ultimate quantity of interest; instead, they serve as intermediate inputs for downstream ecological or statistical estimands. Here, we show that such estimands can be broadly classified into two categories: *perturbation-invariant* estimands, which remain unaffected by PCR bias, and *perturbation-sensitive* estimands, whose values can change substantially under PCR bias. This distinction provides a principled framework for understanding when PCR bias can be safely ignored and when it must be explicitly accounted for. Let ***π***_*dn*_ denote the estimated relative abundance of taxon *d* in the *n*-th sample. A common estimand in differential abundance analyses is the relative log-fold-change: the difference in mean log-ratio abundance between two biological conditions *z*_*n*_ ∈ {0, 1} (e.g., health versus disease):

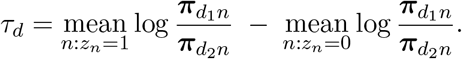

In Corollary 1, we show that *τ*_*d*_ is an example of a broader class of estimands that are *perturbation invariant*. Formally, an estimand *θ* is perturbation invariant if it satisfies

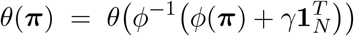

for any *D* − 1-dimensional vector *γ*.

From the perspective of PCR bias, perturbation-invariant estimands are special because they are also invariant to PCR bias. We prove this formally in Theorem 1. This theorem and corollary formalize prior work which hypothesized that relative log-fold-changes were invariant to PCR bias [McLaren et al., 2022, Silverman et al., 2021]. Intuitively, PCR bias, as modeled in Equation (2), acts as a sample-specific perturbation: it shifts all log-ratio coordinates by the term *x*_*n*_***β***. This shift is analogous to adding a global nuisance term 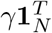, which uniformly affects all log-ratios and therefore cancels out in any estimand based solely on contrasts (i.e., differences between taxa rather than their absolute log-ratio coordinates). Thus, perturbation-invariant estimands are unaffected by PCR bias. In the following section, we turn to perturbation-sensitive estimands, which do not share this invariance and can be strongly influenced by PCR bias.

### 2.3 Common Ecological *α*-Diversity Analyses are Impacted by PCR Bias

The use of *α*-diversity metrics in microbiome research has become ubiquitous with the rise of next-generation sequencing (NGS). These metrics provide a quantitative description of within-sample diversity, capturing aspects of richness and evenness, and are central to ecological and microbiological studies. Reliable and reproducible estimates are particularly important because microbial community structure can strongly influence host physiology and ecosystem stability [Cassol et al., 2025]. However, unlike relative log-fold-change estimands (*τ*_*d*_), we find that *α*-diversity estimands are perturbation sensitive and therefore susceptible to PCR bias.

We evaluated four commonly used *α*-diversity metrics derived from relative abundances: Shannon’s index, Simpson’s index, the Gini coefficient, and the Aitchison norm[Willis and Martin, 2020, Lian et al., 2024, Egozcue and Pawlowsky-Glahn, 2019]. Using the Bayesian multinomial logistic-normal model described in Section 2.1 and data from Silverman et al. [2021], we estimated posterior distributions of these metrics for the true (pre-amplification) compositions and for compositions after 35 PCR cycles (Figure 1). The dataset comprised ten mock communities with controlled proportions of ten bacterial isolates and four human gut microbiota samples from distinct artificial gut systems. Formally, let *f* : 𝕊^*D*^ → ℝ_+_ denote an *α*-diversity metric on the *D*-dimensional simplex. We define PCR-induced bias as:

**Figure 1.**
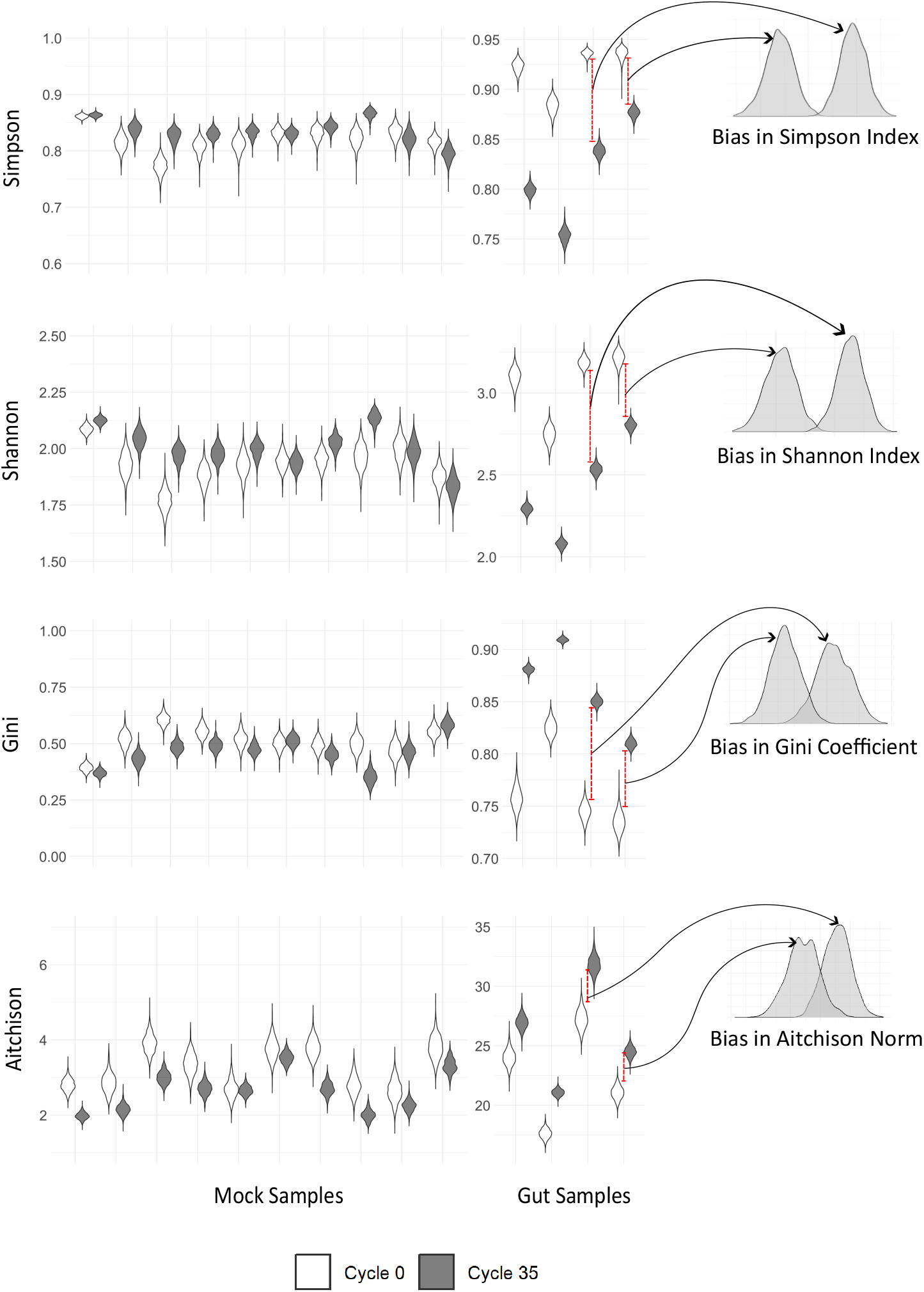
*α*-diversity metrics are impacted by PCR bias. Violin plots show posterior distributions of four *α*-diversity metrics (Shannon, Simpson, Gini, and Aitchison) for *in vitro* and *ex vivo* data, estimated before amplification (0^th^ cycle) and after 35 PCR cycles using the model of Silverman et al. [2021]. Density plots to the right show the posterior distributions of the PCR-induced bias (35-cycle value minus 0-cycle value) for two representative samples. PCR introduces substantial, sample-specific shifts, indicating that comparisons of *α*-diversity across groups may reflect technical artifacts rather than true biological differences.

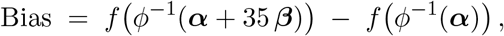

where ***α*** represents the estimated log-ratio composition before amplification (0^th^ cycle) and ***β*** the per-cycle PCR bias.

All four metrics are clearly impacted by PCR bias, with values after 35 PCR cycles substantially differing from their true pre-amplification (0-cycle) values (Figure 1). The magnitude and direction of this bias vary across communities, reflecting its dependence on the underlying composition. Consequently, PCR bias can distort not only absolute estimates of diversity but also comparisons across experimental conditions—what we refer to as differential *α*-diversity analyses, such as studies testing whether *α*-diversity differs between cases and controls (e.g., Van Syoc et al. [2022]).

To assess how strongly PCR bias could affect differential *α*-diversity analyses, we used mock community data from Silverman et al. [2021]. Because the original study did not include natural experimental groupings, we applied an optimization procedure to construct groupings that maximized changes in statistical signal across PCR cycles. We quantified this change as 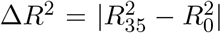, where 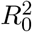 and 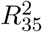 are ANOVA *R*^2^ values at 0 and 35 PCR cycles, respectively. All four metrics showed substantial shifts: Simpson’s index had a Δ*R*^2^ of 0.37, Shannon and Aitchison indices each reached 0.48, and the Gini coefficient exhibited the largest shift at 0.54. These results demonstrate that PCR bias could produce large apparent differences between groups, underscoring the need for caution when interpreting differential *α*-diversity analyses.

To illustrate how PCR bias in *α*-diversity depends on community composition, we used a simplified three-taxon system for visualization (Figure 2 and Supplementary Figure S2). This hypothetical example is not drawn from the mock data but instead illustrates how the magnitude and direction of bias depend on a community’s position in the compositional simplex. Bias is minimal near the edges and vertices, where one or a few taxa dominate, but increases toward the interior, where communities are more even. Intuitively, highly uneven communities are less affected because small changes in relative abundance have little impact on diversity measures: Shannon and Simpson indices are already low in skewed communities, the Gini coefficient is near its upper bound, and the Aitchison norm changes little because extreme compositions occupy fixed positions in log-ratio space. In contrast, even communities are more susceptible—small perturbations can substantially alter the relative balance of taxa, changing Shannon and Simpson diversity, increasing the Gini coefficient, and pushing the Aitchison norm toward more extreme log-ratio coordinates. These visualizations provide intuition for why some communities in the mock and gut data exhibited large shifts in differential *α*-diversity analyses, whereas others were less affected.

**Figure 2.**
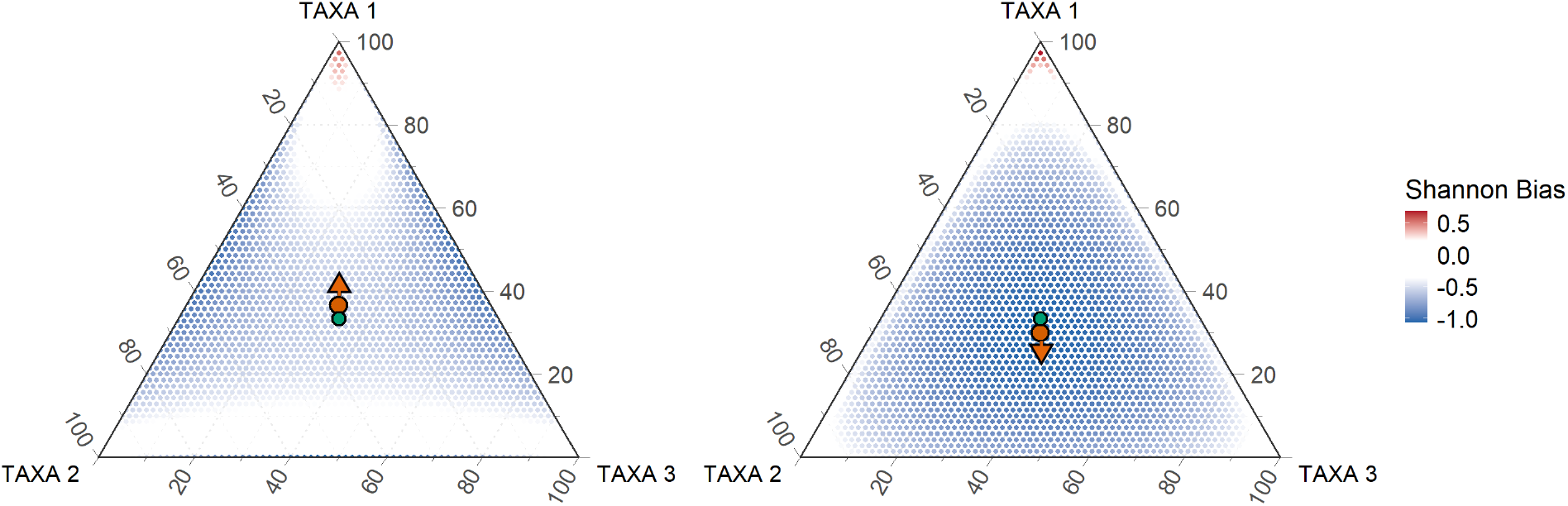
Bias in Shannon index varies across the compositional space. Simplex plots illustrate how PCR amplification alters Shannon diversity under two different bias scenarios. Each point represents an initial (pre-amplification) three-taxon composition, and its color indicates the change in Shannon diversity after 35 PCR cycles. The orange arrow shows the direction and magnitude of the PCR bias vector in composition space, and the green dot marks the unbiased origin. Even small differences in amplification efficiency produce highly non-uniform distortions, with the magnitude of bias depending strongly on the initial community composition.

### 2.4 Ecological *β*-Diversity Analyses are Also Impacted by PCR Bias

*β*-diversity metrics quantify differences in community composition between samples and are widely used in microbiome research when *α*-diversities alone cannot distinguish communities with similar richness and evenness but different taxonomic structures. Among commonly used indices, Weighted-UniFrac incorporates phylogenetic relationships by weighting abundance differences by branch length, whereas Bray-Curtis measures compositional dissimilarity based on relative abundances [Whittaker, 1960, Lozupone et al., 2007]. Because both metrics are perturbation sensitive, they may be affected by PCR bias. Formally, for a *β*-diversity metric *g* : 𝕊^*D*^ × 𝕊^*D*^ → ℝ_+_, we define bias as:

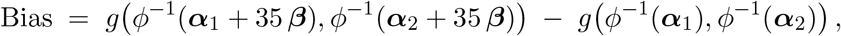

where ***α***_1_ and ***α***_2_ are the log-ratio compositions of the two communities before amplification.

Using the Bayesian multinomial logistic model from Silverman et al. [2021] and the same human gut data described in Section 2.3, we estimated pairwise Bray-Curtis and Weighted-UniFrac distances before amplification (0 cycles) and after 35 PCR cycles (Figure 3). PCR introduced substantial, sample-specific shifts in both metrics, with the magnitude of bias varying considerably across sample pairs.

**Figure 3.**
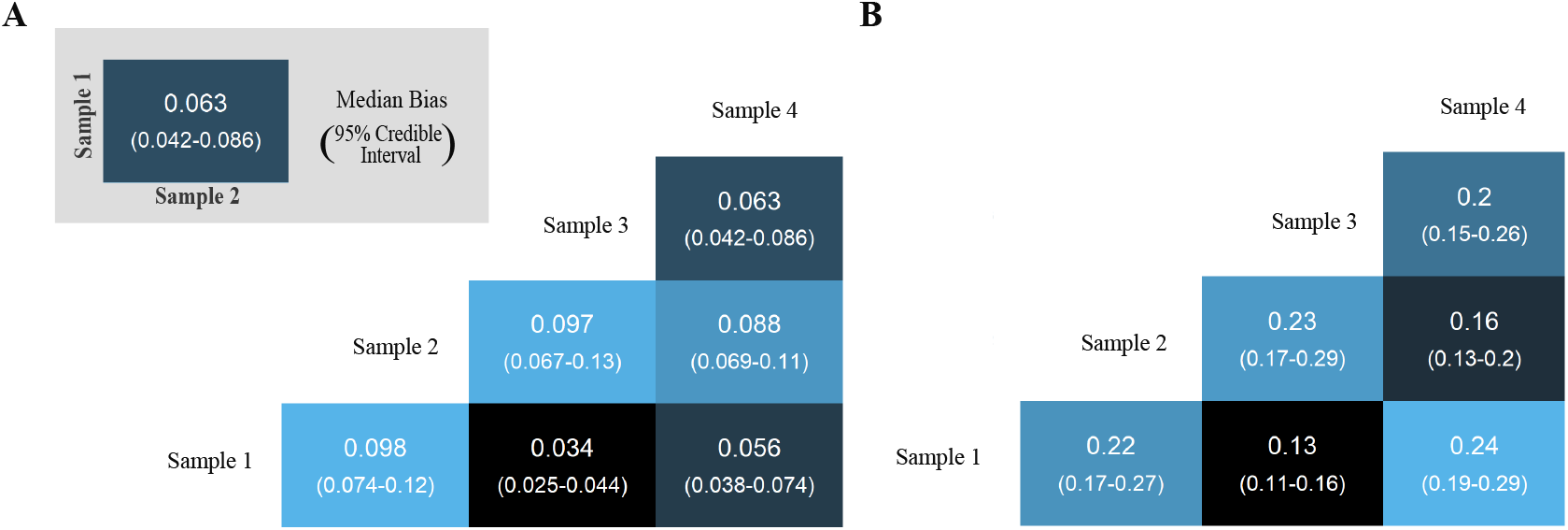
PCR bias alters pairwise *β*-diversity estimates. Heatmaps show the change in Weighted-UniFrac (A) and Bray-Curtis (B) distances between pairs of gut microbiome samples after 35 PCR cycles compared to the pre-amplification (0-cycle) values. Each tile shows the posterior median bias, with 95% credible intervals in parentheses. Variation across pairs indicates that PCR bias introduces systematic, sample-specific distortions in inter-sample dissimilarities.

To evaluate how strongly PCR bias could affect downstream inference, we used the mock community data to test a worst-case scenario for differential *β*-diversity analyses. Because the original study lacked natural experimental groupings, we applied an optimization procedure to construct groupings that maximized the change in PERMANOVA *R*^2^ across PCR cycles. We defined this change as 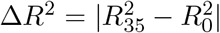, where 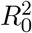 and 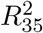 are PERMANOVA *R*^2^ values at 0 and 35 cycles, respectively. The optimized configuration yielded modest but non-negligible shifts: Δ*R*^2^ = 0.08 for Bray-Curtis and 0.12 for Weighted-UniFrac. These results indicate that PCR bias can plausibly distort community-level comparisons.

Finally, to build intuition for why some community pairs are more affected than others, we visualized PCR-induced bias using a simplified three-taxon system (Figure 4 and Supplementary Figure S4). This hypothetical example, included for visualization only, shows how bias depends on the positions of the two communities in the compositional simplex. Bias is minimal when the PCR-induced compositional shift is approximately orthogonal to the differences between the two communities, producing a characteristic X-shaped region of low bias in the simplex. Conversely, when PCR bias aligns with the key taxonomic or phylogenetic differences between communities, even small amplification shifts can substantially inflate or deflate *β*-diversity estimates. This provides intuition for the heterogeneity in bias observed in the mock and gut data.

**Figure 4.**
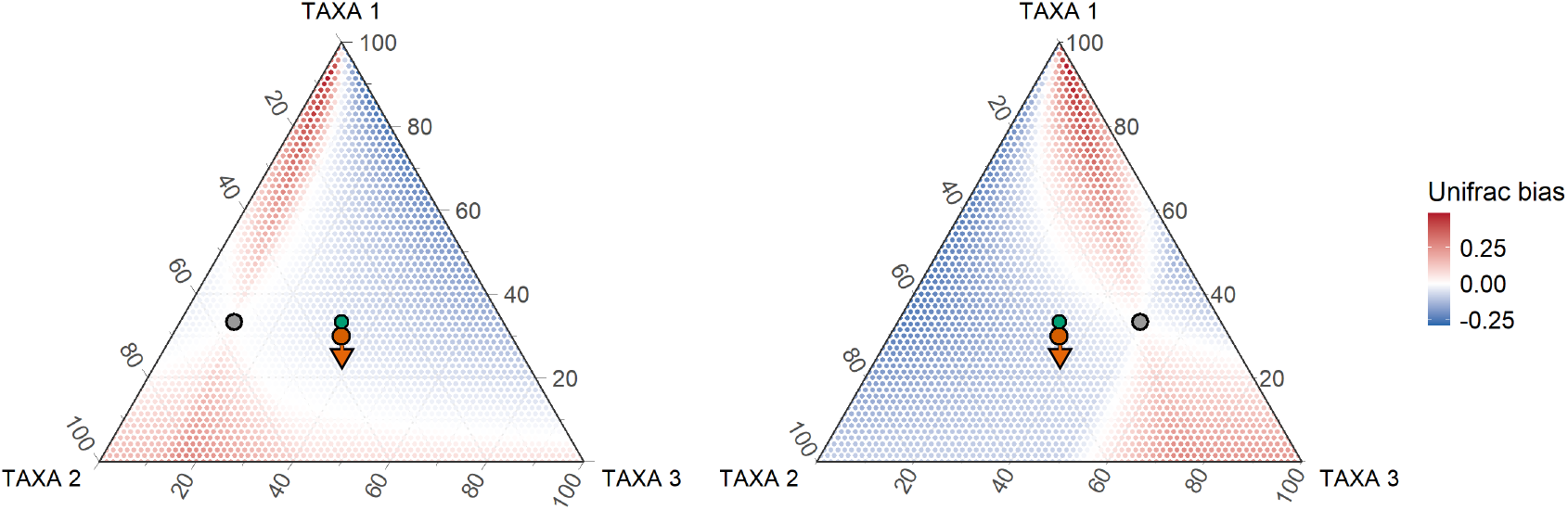
Bias in Weighted-UniFrac depends on community composition. Simplex plots show how Weighted-UniFrac bias varies as the second community (*x*_2_) moves across the compositional simplex relative to a fixed reference (*x*_1_), with PCR bias held constant. Color indicates the change in Weighted-UniFrac (35-cycle minus 0-cycle value). The orange arrow shows the direction of PCR bias, the green dot marks the unbiased origin, and the grey circle marks the fixed reference. Regions of minimal bias form a characteristic X-shaped band, illustrating that bias is lowest when PCR-induced shifts are orthogonal to the differences between communities.

### 2.5 Aitchison Distance Remains Invariant to PCR Bias

Unlike other *β*-diversity metrics, the Aitchison distance is unaffected by PCR bias because it is perturbation invariant, meaning it is unchanged by the additive shifts in log-ratio space introduced during amplification. For two communities *n*_1_ and *n*_2_, the Aitchison distance is defined as:

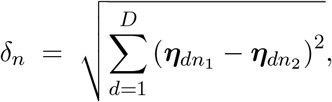

where ***η***_*n*_ = *ϕ*(***π***_*n*_) is the log-ratio transformed relative abundance vector for sample *n*. Intuitively, this distance measures how far apart two communities are in log-ratio space: a value of zero indicates identical relative compositions, whereas larger values reflect greater differences in the compositional structure of the communities. Because the distance is Euclidean, differences accumulate additively across log-ratios, making it straightforward to interpret which log-ratios contribute most to the separation.

As we prove in Corollary 2, the additive perturbations introduced by PCR amplification cancel in this log-ratio representation, leaving the distance unchanged. This invariance has important practical implications. Unlike Bray-Curtis or Weighted UniFrac, which we show to be highly sensitive to PCR bias, the Aitchison distance provides a robust framework for *β*-diversity analysis. Because it remains stable even when amplification bias is present, the Aitchison distance helps ensure that observed differences between communities reflect genuine ecological or experimental variation rather than PCR-induced artifacts.

## 3 Discussion

While PCR is a routine step in microbiome profiling, differences in taxon-specific amplification efficiencies can introduce substantial bias in estimated community compositions. Primer mismatch effects occur during the initial amplification cycles, but multiple studies have shown that non–primer-mismatch (NPM) sources of bias dominate PCR bias overall [Gimpel et al., 2024, Korvigo et al., 2022, Silverman et al., 2021]. Here we extended prior studies of NPM sources of PCR bias (called PCR bias for brevity) to systematically evaluate how these biases affect ecological diversity analyses.

Our results prove that PCR bias alters both *α*- and *β*-diversity estimates, with the magnitude and direction of distortion depending strongly on the underlying community composition. For *α*-diversity, bias is greatest in communities with even taxonomic structures, where small shifts in relative abundances substantially change diversity metrics, whereas highly skewed communities show minimal distortion. For *β*-diversity, bias is lowest when the PCR-induced perturbation is largely orthogonal to the differences between communities, but can be substantial when it aligns with the primary axes of taxonomic or phylogenetic variation. These composition-dependent effects mean that PCR bias can obscure or exaggerate apparent ecological differences between groups. Using ANOVA for *α*-diversities and PERMANOVA for *β*-diversities, we further demonstrate that bias can amplify or suppress group-level signals, potentially leading to misinterpretation in ordination, clustering, or differential diversity analyses.

On the other hand, when all samples are amplified under the same PCR protocol, the multiplicative model of PCR bias implies that taxon-specific efficiencies are applied uniformly, producing a constant additive shift in log-ratio space. As we prove formally, perturbation-invariant estimands, such as differential log-ratios and the Aitchison distance, are unaffected by such shifts, making them well-suited for downstream analyses when consistent amplification protocols are used.

Given these findings, we recommend using compositional metrics based on log-ratio transformations, such as the Aitchison distance, whenever possible for analyses of PCR-amplified microbiome data. These methods are perturbation invariant and therefore unaffected by PCR bias. However, the choice of metric should ultimately be guided by the scientific question, and some analyses may require traditional diversity measures (e.g., Shannon, Simpson, Gini, Bray-Curtis, Weighted UniFrac). Because these metrics are sensitive to PCR bias and the magnitude of bias depends on community composition, they should be interpreted with caution—both in terms of their absolute values and when used for between-sample comparisons. When such traditional metrics are necessary, we recommend performing calibration experiments (e.g., sequencing the same community with different numbers of PCR cycles) to estimate community-specific amplification biases and adjust the diversity measures following techniques introduced in this article and in Silverman et al. [2021].

Our analysis has focused on continuous functions of relative abundances and has not considered presence/absence-based metrics (e.g., Jaccard, unweighted UniFrac), which are more sensitive to detection thresholds and sequencing noise. Although PCR bias can theoretically push rare taxa below detection, modeling such effects requires explicitly accounting for detection variability and is beyond the scope of this study. Future work should assess how PCR bias interacts with presence/absence metrics and evaluate its impact across a broader range of experimental settings. Independent validation data, such as technical replicates or mock communities, will be essential for determining the operational relevance of these findings in real-world microbiome studies. As more datasets become available, our understanding of the ecological consequences of PCR bias will continue to improve.

## 4 Methods

### 4.1 PCR Bias Model with Covariates

We modeled PCR bias using the Bayesian multinomial logistic-normal framework introduced by Silverman et al. [2021], which extends the standard exponential amplification model to accommodate multiple taxa, technical noise, and structured sample covariates. The model is specified as:

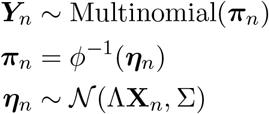

where *Y*_*n*_ is the vector of sequencing read counts for sample *n, π*_*n*_ is the corresponding vector of relative abundances, and *η*_*n*_ = *ϕ*(*π*_*n*_) denotes the log-ratio transformed abundance. The matrix Λ ∈ ℝ^(*D−*1)*×p*^ contains regression coefficients, Σ ∈ ℝ^(*D−*1)*×*(*D−*1)^ is the residual covariance, and *X*_*n*_ ∈ ℝ^*p*^ is a covariate vector specifying sample-level features.

This formulation allows for flexible modeling of complex experimental designs. For example, to jointly model samples from two biological communities with varying numbers of PCR cycles, we can construct a design matrix *X* ∈ ℝ^*p×N*^, where each column *X*_*n*_ includes a one-hot encoding of community membership and a numeric variable for PCR cycle count:

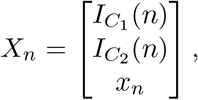

with *I*_*C*_*i* (*n*) = 1 if sample *n* belongs to community *C*_*i*_ (and zero otherwise), and *x*_*n*_ denoting the number of PCR cycles applied. In this case, the coefficient matrix Λ takes the form:

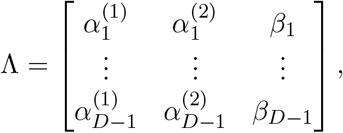

where 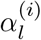 is the baseline (cycle-0) log-ratio abundance of the *l*^th^ contrast in community *C*_*i*_, and *β*_*l*_ represents the per-cycle PCR amplification bias on that log-ratio. This structure enables simultaneous inference of both biological variation and amplification bias, even when only some samples vary in PCR cycle number.

All model fitting was performed in R using the fido package Silverman et al. [2022].

### 4.2 Datasets and Preprocessing

We analyzed two publicly available datasets originally described by Silverman et al. [2021]: ten in vitro mock communities composed of known proportions of ten bacterial isolates and four ex vivo human gut microbiota samples derived from distinct artificial gut systems. Each sample was split into three replicates, with each replicate subjected to a different number of PCR cycles prior to sequencing, enabling estimation of all model parameters. All analyses were conducted using the same preprocessed datasets and parameter settings as in the original study. We re-ran the publicly available fido pipeline without modification to estimate posterior distributions of *α*_*n*_ (true relative abundances for each community at cycle 0) and *β* (taxon-specific relative amplification efficiencies).

### 4.3 Diversity Metrics and Bias Estimation

We evaluated the effect of PCR bias on commonly used ecological diversity measures. Posterior samples from the fitted model were used to estimate taxon relative abundances at 0 and 35 PCR cycles, which served as inputs for downstream calculations.

#### *α*-diversity

We analyzed four *α*-diversity metrics: Shannon index, Simpson index, Gini coefficient, and the Aitchison norm. Shannon, Simpson, and Gini indices were computed using custom R functions. The Aitchison norm was calculated as the Euclidean norm of centered log-ratio (CLR) transformed abundance vectors. PCR-induced bias for each metric was defined as:

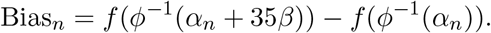

#### *β*-diversity

We evaluated three *β*-diversity metrics: Bray-Curtis dissimilarity, Weighted UniFrac, and Aitchison distance. Bray-Curtis dissimilarities were calculated using the `vegdist()` function in the `vegan` package. Weighted `UniFrac` distances were computed using the `UniFrac()` function in the `phyloseq` package with a pruned `greengenes2` phylogenetic tree. Aitchison distances were calculated as Euclidean distances in Centered Log-Ratio (CLR) space. Bias was defined analogously to *α*-diversity metrics:

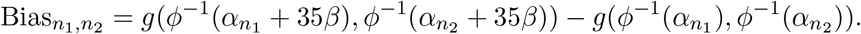

### 4.4 Statistical Analyses

#### Alpha diversity

To quantify the effect of PCR bias on group-level diversity comparisons, we computed mean diversity values per sample at both cycle 0 and cycle 35 and evaluated group differences using ANOVA. A genetic algorithm was applied to identify binary groupings of samples that maximized the absolute change in ANOVA *R*^2^ across PCR cycles:

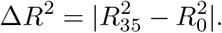

#### Beta diversity

To assess changes in community structure, we performed PERMANOVA (`adonis2() in vegan`) on each *β*-diversity distance matrix. Particle swarm optimization (`psoptim() in R`) was used to find the grouping of samples that maximized changes in PERMANOVA *R*^2^ across PCR cycles, providing a worst-case estimate of how much amplification could distort community-level comparisons.

### 4.5 Visualization of Composition-Dependent Bias

To build intuition for composition-dependent effects, we simulated PCR bias in a simplified three-taxon system. Bias was evaluated across a grid of compositions in the 3-part simplex, holding the amplification bias vector constant. This visualization highlights how bias depends on the position of communities in the simplex and explains variability observed in the mock community analyses.

## 4.6 Data and Code Avaibility

All code and data used in this study are available at https://github.com/dharmikrathod/pcr_bias_code.

## Acknowledgments

The authors would like to thank Dr. Rachel Silverman for her manuscript comments. JDS was supported by NIGMS R01GM148972-01.

## APPENDIX A Supplementary Proofs

**Theorem 1** (Perturbation-Invariant Estimands are Invariant to PCR Bias). *Let* Ψ ∈ ℝ^(*D−*1)*×D*^ *be a rank D* −1 *contrast matrix with columns summing to zero, and suppose the observed compositional vector w*_*n*_ ∈ *S*^*D*^ *for sample n follows the linear log-contrast model from eq*. (2):

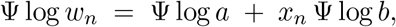

*where a* ∈ *S*^*D*^ *is the true (pre-amplification) composition and b* ∈ ℝ^*D*^ *is the vector of taxon-specific PCR efficiencies*.

*Let θ*(·) *be any estimand that is perturbation invariant, i*.*e*.,

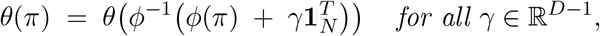

*where π* ∈ *S*^*D*^ *is any compositional vector, ϕ*(·) *is a log-ratio transform, and ϕ*^*−*1^(·) *its inverse*.

*Then θ is invariant to PCR bias:*

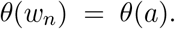

*Proof*. From the PCR bias model:

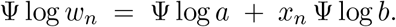

Define the log-ratio transformed vectors:

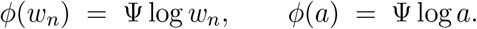

Then PCR bias appears as an additive perturbation in log-ratio space:

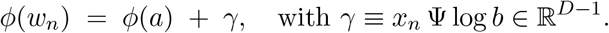

By the perturbation-invariance property of *θ*:

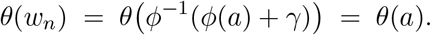

Thus, *θ* is unaffected by PCR bias under this model.

**Corollary 1** (Differential Log-Ratio Abundance is Invariant to PCR Bias). *Let the true differential log-ratio abundance of two taxa between two biological conditions z*_*n*_ ∈ {0, 1} *be defined as:*

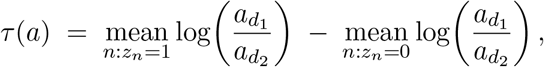

*where a* ∈ *S*^*D*^ *is the true (pre-amplification) composition. Suppose the observed composition w*_*n*_ ∈ *S*^*D*^ *follows the PCR bias model from Equation* (2):

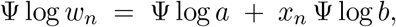

*with b* ∈ ℝ^*D*^ *the vector of taxon-specific PCR efficiencies and x*_*n*_ *the number of PCR cycles*.

*Then the differential log-ratio abundance is invariant to PCR bias:*

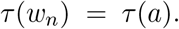

*Proof*. The estimand *τ* depends only on differences in log-ratios of two taxa (*d*_1_ and *d*_2_) between groups. Adding a constant perturbation *γ* to the log-ratio space shifts all coordinates by the same amount, but such a constant cancels when computing log-ratio differences and group means. Formally:

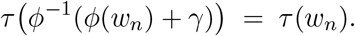

Thus, *τ* satisfies the perturbation-invariance condition of Theorem 1. Applying the theorem:

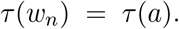

**Corollary 2** (Aitchison Distance is Invariant to PCR Bias). *Let the true Aitchison distance between two communities n*_1_ *and n*_2_ *before PCR amplification be:*

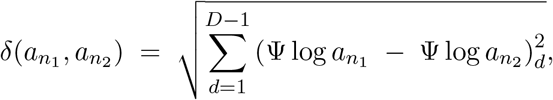

*where a*_*n*_ ∈ *S*^*D*^ *is the true (pre-amplification) composition and* Ψ *is a rank-*(*D* − 1) *contrast matrix with columns summing to zero. Suppose the observed composition w*_*n*_ ∈ *S*^*D*^ *follows the PCR bias model from Equation* (2):

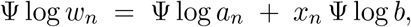

*with b* ∈ ℝ^*D*^ *the vector of taxon-specific PCR efficiencies and x*_*n*_ *the number of PCR cycles*.

*Then the Aitchison distance is invariant to PCR bias:*

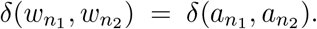

*Proof*. The Aitchison distance depends only on pairwise differences in log-ratio coordinates:

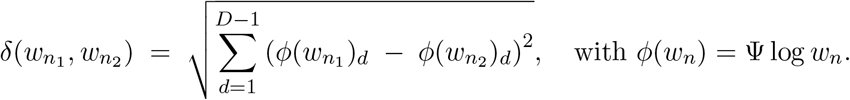

Adding a constant perturbation *γ* shifts all log-ratio coordinates by the same amount, but such a constant cancels when taking differences:

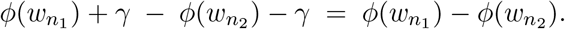

Thus:

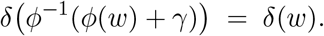

The Aitchison distance therefore satisfies the perturbation-invariance condition of Theorem 1. Applying the theorem:

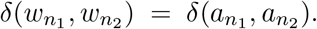

## Appendix B B Supplementary Figures and Tables

**Figure S1.**
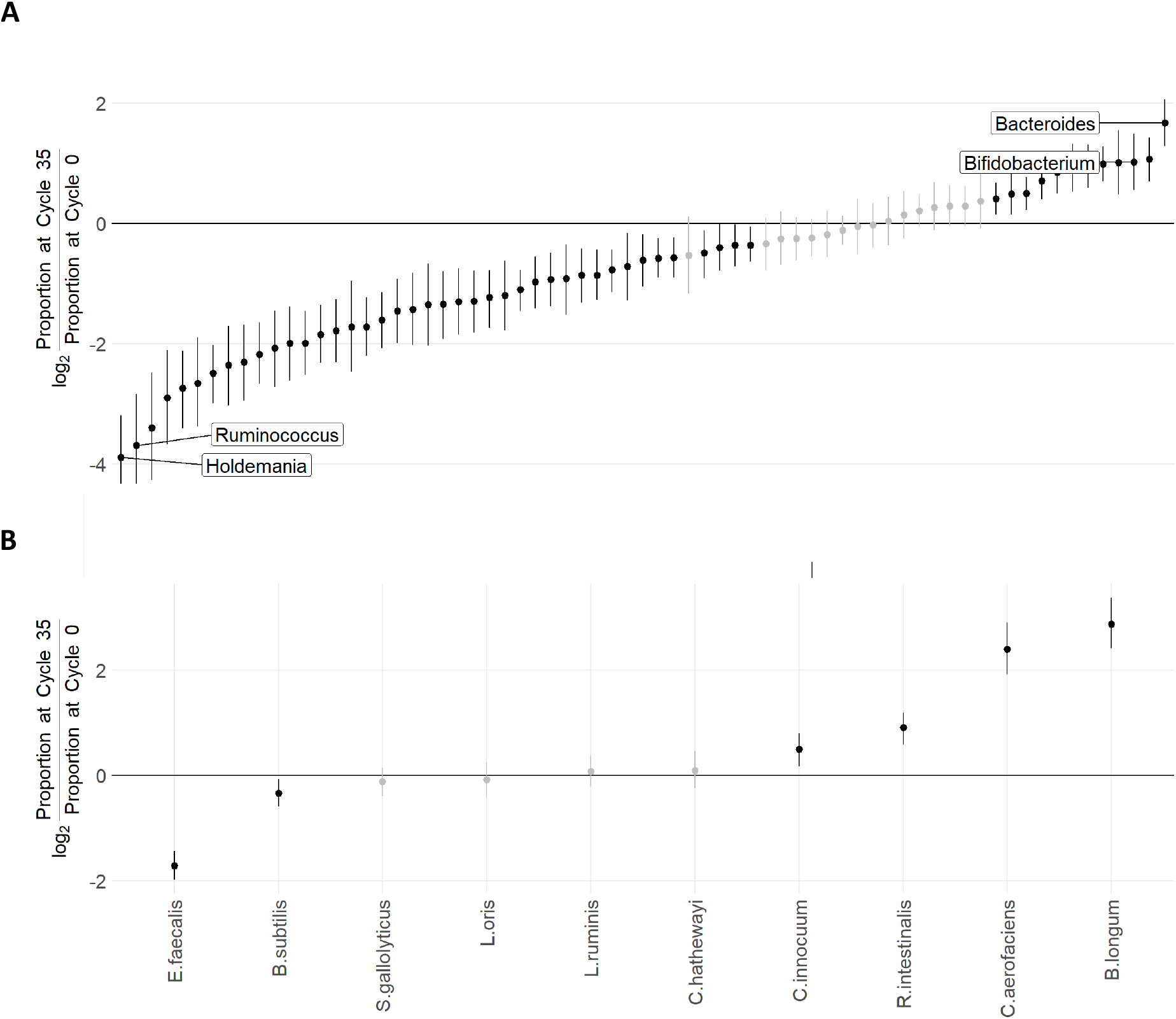
PCR bias in human gut and mock communities. Bias was quantified as the log_2_- ratio of each taxon’s estimated relative abundance after 35 PCR cycles (cycle 35) to its estimated pre-amplification abundance (cycle 0) for (A) human gut microbiota and (B) mock communities from Silverman et al. [2021]. A log_2_ value of 2 indicates a fourfold overestimation due to PCR, whereas -2 indicates a fourfold underestimation. Points show posterior medians, and bars denote 95% credible intervals. Taxa with credible intervals excluding zero (statistically credible bias) are highlighted in black.

**Figure S2.**
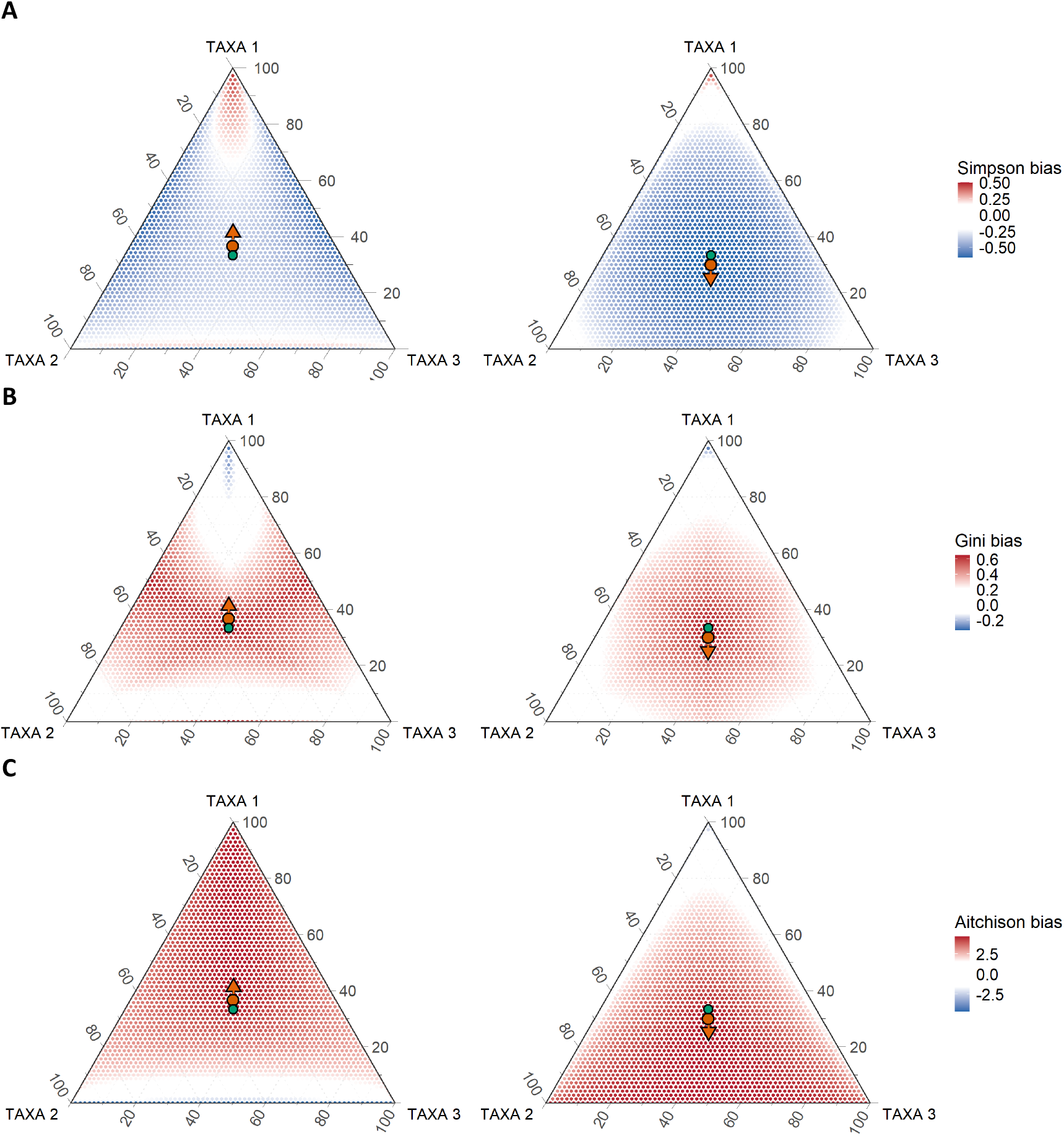
Composition-dependent bias in Simpson, Gini, and Aitchison diversity. Simplex plots show PCR-induced changes in (A) Simpson diversity, (B) Gini coefficient, and (C) Aitchison norm after 35 cycles under two different bias vectors (orange arrow). Each point represents an initial pre-amplification composition in a three-part simplex, colored by the change in the corresponding metric after PCR. The green dot marks the unbiased origin, and the orange arrow indicates the direction of the bias vector. Even small differences in amplification efficiency produce highly non-uniform distortions, with bias magnitude varying systematically across the compositional space.

**Figure S3.**
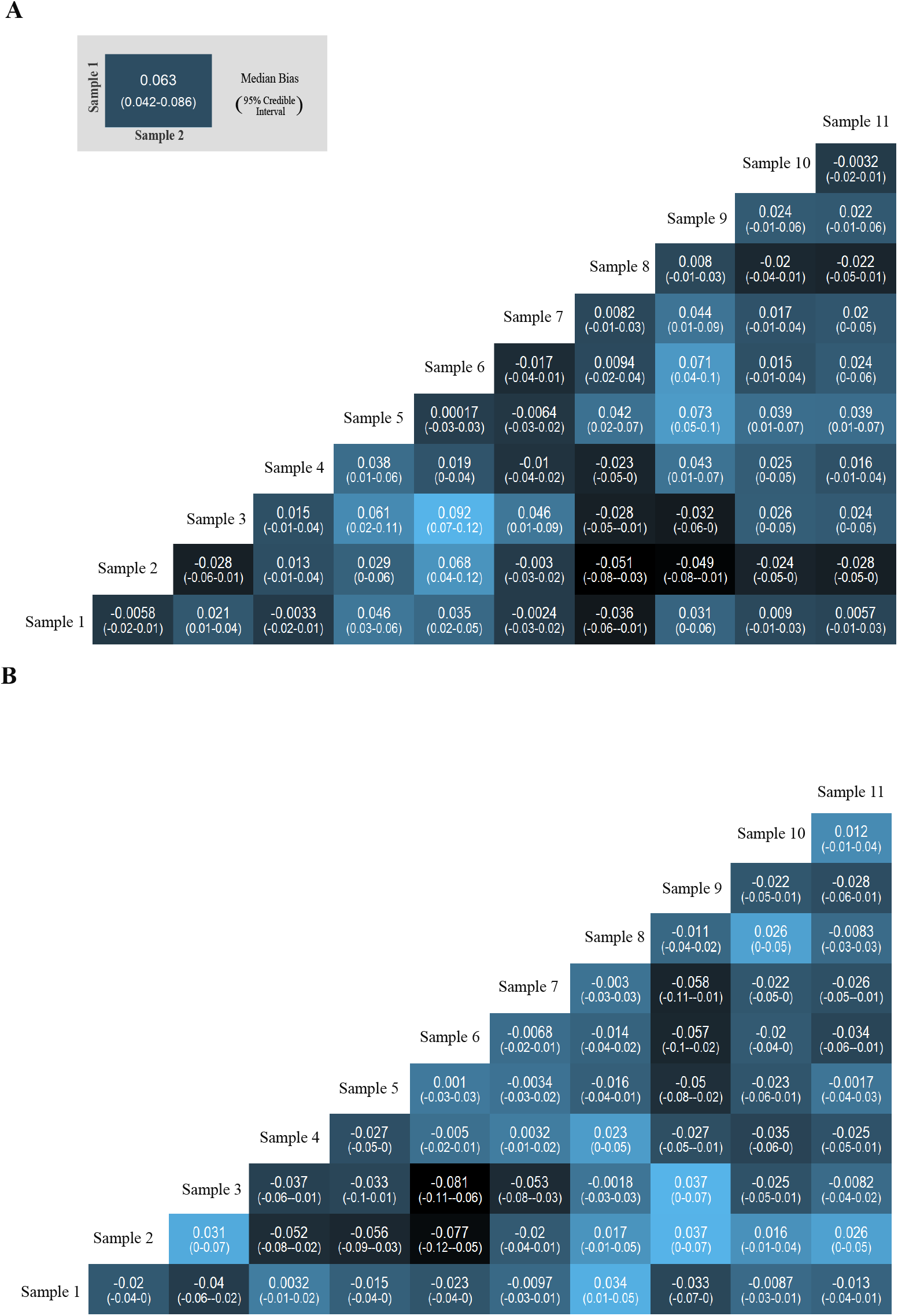
Pairwise *β*-diversity bias in mock communities. Heatmaps show changes in (A) Weighted UniFrac and (B) Bray-Curtis distances between pairs of mock community samples after 35 PCR cycles relative to their pre-amplification (cycle 0) values. Each tile shows the posterior median change, with 95% credible intervals in parentheses. Rows and columns correspond to sample indices. Variation in bias across pairs indicates that PCR amplification introduces systematic and sample-specific distortions in inter-sample dissimilarities.

**Figure S4.**
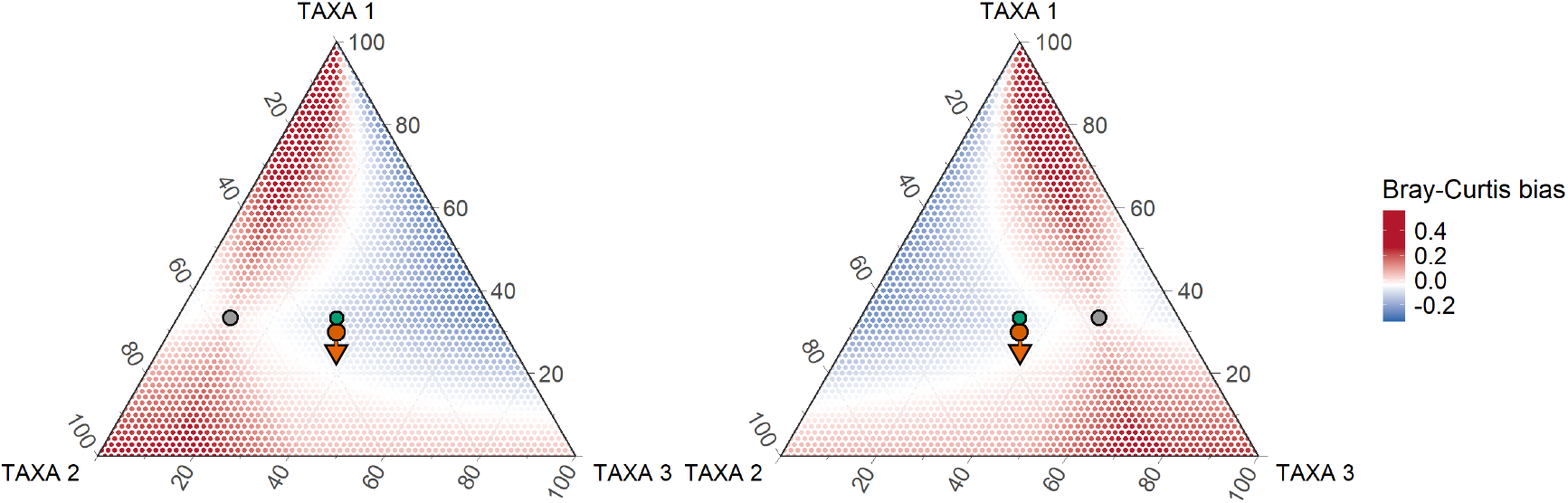
Composition-dependent bias in Bray-Curtis dissimilarity. Change in Bray-Curtis distance is shown as the second community (*x*_2_) varies across the simplex relative to a fixed reference composition (*x*_1_), with PCR bias held constant. Colors indicate the difference between the Bray-Curtis value at 35 PCR cycles and its true pre-amplification value. The green dot marks the unbiased origin, the orange arrow indicates the direction of PCR bias, and the grey circle marks the fixed reference community (*x*_1_). Distortion depends strongly on the initial compositions of both communities, with bias magnitude varying systematically across the simplex.

